# Mucin methacryloyl (MuMA) based hydrogels: tunable features for tissue engineering and drug release control

**DOI:** 10.64898/2026.11.22.768418

**Authors:** Enam Khalil, Enas Alkhafaji, Christian Umemura, Reihaneh Haghniaz, Unnati Modi, Satoru Kawakita, ALSayed Sallam, Fatemeh Zethabi, Neda Farhadi, Mai Jaber, Behareh Mir, Anindita Sarkar, Muhammad Al Zweiri, Samsuzzoha Mondal, Rajesh Vasita, Kalpana Mandal

## Abstract

Biopolymer-based hydrogels are widely used in tissue engineering and controlled drug delivery; however, simultaneously achieving tunable mechanical properties, favorable pore architecture, and controlled release characteristics remains challenging. In this study, we developed hybrid hydrogels based on mucin methacryloyl (MuMA), a methacrylated derivative of the naturally abundant glycoprotein mucin, combined with gelatin methacryloyl (GelMA) and glycerin to create materials with tailored physicochemical and biological properties.

Compared with MuMA-only hydrogels, MuMA-GelMA hybrid hydrogels exhibited tunable mechanical behavior, larger pore sizes, and lower phase shift values (Gʺ/Gʹ), indicating enhanced elastic characteristics. These structural and rheological properties supported improved cell adhesion and proliferation while facilitating nutrient transport within softer matrices. The hybrid hydrogels also demonstrated slower drug release kinetics, excellent biocompatibility, and enhanced water retention, highlighting their potential for cell-laden tissue constructs and sustained drug delivery applications.

Furthermore, incorporation of glycerin as a plasticizing agent produced flexible and resilient hydrogel films with improved handling characteristics. This added functionality expands the utility of MuMA-based materials for the development of advanced biomedical devices. Owing to the unique chemical structure of mucin, MuMA readily participates in both physical and chemical interactions with polymers and therapeutic agents, providing a versatile platform for engineering hydrogels with tunable properties for tissue regeneration and controlled drug delivery.

## INTRODUCTION

Hydrogels are three-dimensional networks of hydrophilic polymers formed through physical or chemical crosslinking. Upon crosslinking, the polymers form a water-insoluble network capable of absorbing and retaining large amounts of water while maintaining their structural integrity. The network structure and the swelling profile can be tailored according to intended application (Todd R. Hoare a 2008). Hydrogels have several applications in many fields including tissue engineering (Su Ryon Shin 2013) (Natan R. Barros 2025) and drug delivery (Zhimin Luo and Khademhosseini 2018). Natural and synthetic polymers are used in the production of hydrogels (Su Ryon Shin 2016) (Su Ryon Shin 2013) (Nguyen Hoc Thang 2023) (Song Chen 2023). Natural polymer-based hydrogels are preferred in biomedical applications due to their similarity to the extracellular matrix (ECM) (Natan R. Barros 2021). As drug carriers; hydrogels can be designed for specific delivery routes (Jianyu Li 2016). Natural polymers-based hydrogels like gelatin, alginate, chitosan and mucin cover a wide range of physicochemical properties (Su Ryon Shin 2016) (Nguyen Hoc Thang 2023). In the last ten years, mucin-based hydrogels show increasing interest for different applications including osteoimmunomodulation (Song Chen 2023), drug delivery (Duffy 2015) and the field of tissue reconstruction and repair (Elena Olaret 2021).

Mucins are the key functional components of the mucosal layer, which protect the underlying epithelial tissues (Katherine Joyner 2019). Mucins secreted from different organs, such as the colon, gut, or salivary glands, coat and protect the epithelial lining of the organs. Transmembrane mucins participate in cellular signaling pathways, which are altered during many diseases, including cancer, by overexpressing mucins (Kufe 2009).

There are two commercially available highly investigated mucin types: porcine gastric mucin (PGM) (Bradley S. Turner 1995, JanaT.Reh 2025, Mar Collado-González 2025), and bovine submaxillary mucin (BSM) (Duffy 2015, Hanna Rulff and Marcus A. Mall 2024). However, these mucins do not form hydrogels, presumably because of their purification process (Veronika J. Schömig 2016) (Katherine Joyner 2019). PGM has been observed to aggregate in aqueous solutions resulting from a lower percentage of negatively charged amino acids, forming less elastic and more viscous films. In aqueous solutions, PGM forms thin loosely bonded films on a hydrophobic Polydimethylsiloxane (PDMS) surface, the loose structure of these films was related to the localized distribution of negative charges at the terminal regions and the hydrophobic patches being buried within the globular conformation of the PGM (Gleb E. Yakubov 2007, Emanuela Di Cola 2008, Jan Busk Madsen 2016). Mucin structure contains different functional groups that can be engaged in hydrogel formation. The presence of a hydroxyl group in the terminal segment of mucin renders it capable of forming many hydrogen bonds that interact with a variety of polymers. Moreover, due to the presence of cysteine-rich regions, mucins can easily participate in intermolecular crosslinking by forming inter- and intra-molecular disulfide bonds resulting in mesh-like structure formation (Katherine Joyner 2019). In addition, Mucins are resistant to proteolysis and enhance the ability to hold water due to their glycosylated properties (Veronika J. Schömig 2016). Mucin hydrogels were generated using different cross-linking methods including tetrazine- and norbornene-mediated protein-protein crosslinking, UV cross-linking (Duffy 2015, Wang 2025) and thiol-based disulfide bridge formation (Katherine Joyner 2019), which have resulted in viscoelastic properties comparable to native mucus. Other methods cross linkers were also used including tannic acid (Chiara Gunnella 2025, Yilin Yue 2026), boric acid (Yuki Nakamura 2026) and genipin, (Mar Collado-González 2025).

Mucin hydrogels are attracting increasing attention for various biomedical applications including immune activity (Kun Jiang 2021) biomedical coatings (Tai Peng 2023) and wound healing (Serafim 2018). Mucin-based hydrogels also act as multifunctional biomaterials owing to their excellent biocompatibility, high water-retention capacity, biodegradability, and ability to recapitulate several structural and biological features of the native extracellular matrix. These properties have enabled their application in a wide range of biomedical fields, including immunomodulation, tissue engineering, and drug delivery (Crouzier et al., 2018; Yan et al., 2020). Unlike conventional hydrogels, mucin hydrogels actively regulate the host immune response by modulating macrophage activation and suppressing the foreign body response, thereby creating a favorable microenvironment for tissue regeneration (Yan et al., 2020; Crouzier et al., 2022). Furthermore, hybrid 3D scaffolds composed of mucin and collagen have demonstrated significantly enhanced osteogenic differentiation and bone regeneration compared with scaffolds containing either polymer alone, highlighting the synergistic interaction between collagen and mucin in promoting tissue repair (Shi et al., 2023) (Barik, Bejugam et al. 2021). In addition to their structural role, the amphoteric nature of mucins, arising from the coexistence of positively and negatively charged domains within their heavily glycosylated structure, enables electrostatic interactions with a broad spectrum of therapeutic molecules. Consequently, mucin-based hydrogels are promising carriers for the encapsulation and controlled release of drugs with diverse physicochemical properties, including small molecules, proteins, peptides, and nucleic acids, making them potential candidate as carriers for drugs with different physicochemical properties (Turner 2018) (Bansil and Turner, 2018; Crouzier et al., 2022).

In general, hydrogel-behavior can be conveniently tailored for any of their wide application using suitable additives (Hyeongho Shin 2012) (Hyeongho Shin 2012, Su Ryon Shin 2012). In tissue engineering, the cellular microenvironment represented by the ECM, plays a crucial role in cell growth, proliferation, migration, and metastasis (Mandal, Wang et al. 2014, Mandal, Raz-Ben Aroush et al. 2019) (Mora-Boza, Lopez-Ruiz et al. 2020). ECM is comprised of various proteins, such as collagen, fibronectin, glycosaminoglycans, gelatin, and mucin (Ushida, Sato et al. 2022) and it is required to design a suitable biomaterial that mimic in vivo mechanical or biochemical properties (Du, Lo et al. 2008, Mandal, Gong et al. 2020). Various natural and synthetic biopolymers have been explored to create proper cellular microenvironments in the past several years, including mucin (Duffy 2015) or GelMA-based hydrogels (Kan Yue 2015). GelMA hydrogels are widely used and well characterized for their biomedical applications (Kan Yue 2015) (Ashlyn T. Young 2020) (Ewa Walejewska 2024).

The addition of GelMA to mucin based hydroels is expected to enhance the cell adhesion properties of MuMA hydrogel, and to modify its viscoelastic behavior via the expected physical cross linking and interpenetrating network formation between the two polymers (Manoj N.R. 2002).

Using a second interpenetrating polymer is expected to modify the rate of the gelation process and, consequently, the hydrogel’s mechanical properties (Mora-Boza, Lopez-Ruiz et al. 2020, Valeria Nele 2020) where the fast gelation was found to result in uneven crosslinking, leading to cell elongation (Nicole J. Darling 2016). A mucin hydrogel was also modified for controlled curcumin release using pectin (Islam, Wong et al. 2022) while the viscoelastic properties of mucin hydrogels were modified by the addition of salts (Rulff, Schmidt et al. 2024). Plasticizers (glycerin, sorbitol, polyethylene glycol,..) are one of the widely used additives especially when the hydrogels are intended to be used in a sheet or film form (Maram Diab 2021) (Guohou Sun 2018). Plasticizers decrease the polymer glass transition temperature (Tg) that is when the polymer structure turns viscous or rubbery (Maram Diab 2021). At temperatures below the Tg, the polymeric film shows the properties of the amorphous glassy state (i.e., brittleness, stiffness, and fragility) (Soad A Mohamad 2020) which makes the film challenging to handle. Plasticizers decrease the polymer Tg value by being inserted between the polymer chains and splitting them apart, thus increasing the free volume in the hydrogel structure and weakening polymer chains’ mutual interactions. Glycerin was used in the preparation of gelatin hydrogel sheets intended for in utero tissue engineering application. The produced sheets were biocompatible, easy to be rolled up for insertion into the curved target site (Miho Watanabe 2020) (Albert Rosado 2024).

In this work; MuMA based hydrogels were prepared either alone or in presence of additives and investigated for tissue engineering and drug release applications. Gelma was added to improve the hydrogel bioadhesion and to modify its mechanical and swelling properties while glycerin was added to increase flexibility of the produced hydrogel films.

## MATERIALS AND METHODS

### MuMA synthesis and structural characterization

Mucin methacryloylation method was adapted from Duffy *et al* (Duffy 2015) with modifications. One gram powder of PGM type II (Sigma-Aldrich) was suspended in deionized distilled water (DiDW) and stirred at RT for two hours to produce colloidal mucin dispersion with a pH of 4.5. Mucin dispersion was then transferred to an ice bath and placed over a magnetic stirrer. The dispersion pH was adjusted to 8.0 using 2 N NaOH (Fischer Chemical, USA). Then, methacrylic anhydride (MA) (Sigma-Aldrich) was added dropwise to the colloidal mucin dispersion as 1% (v/v). The solution pH was measured and readjusted to pH 8.0 using 2 N NaOH before the next MA addition. The MA addition process was extended for about three hours, and the pH was adjusted for the next five hours, after which changes in the pH became minimal. The reaction vessel was kept at 4 ^0^C overnight. After 24 hours, the pH was assessed, and the reaction vessel was further stirred at 4 ^0^C for another three hours to ensure the completion of the reaction. At this point, the mixture was turbid due to the presence of insoluble mucin components. Next, the dispersion was centrifuged for one hour at 14,000 rpm (Sorvall Legend X1R, ThermoFisher, Germany) at RT. The supernatant was then filtered using a 0.22 μm filtration unit to obtain a clear solution, which was placed in dialysis tubing MWCO 12-14 kD (Spectra/Por molecular porous membrane tubing, Spectrum Laboratories, Inc USA). The dialysis process was performed against 5.0 L DiDW under stirring in an ice bath. The water in the external compartment was changed three times per day for three days. The resulting MuMA solution was frozen at -80 ^0^C and freeze-dried before being stored at -20 ^0^C for further use.

#### NMR characterization

All freeze-dried MuMA samples, were dissolved in dimethyl sulfoxide-d6 (Deuterated DMSO) and analyzed at RT using a Bruker ARX400 NMR.The freeze-dried MuMA samples (20 mg) were partially dissolved in 750 µl DMSO. Then, the MuMA solution was transferred to NMR tubes for analysis. The 1H-NMR chemical shifts were registered in parts per million (δ) for 64 scans. In addition, the following samples were also characterized: crosslinked MuMA (10% MuMA, 1% LAP, crosslinked for 10 minutes using 10 mW/cm^2^ UV energy (at 360 nm)), physical mixture of 10% MuMA with 1% w/w of the photoinitiator, LAP (Lithium phenyl-2,4,6-trimethylbenzoylphosphinate, Sigma-Aldrich, ≥ 95%) and PGM powder.

### Hydrogel preparation

MuMA based hydrogel was prepared by dissolving the MuMA, and 1% (w/v) LAP in phosphate buffer saline (PBS), followed by crosslinking using UV with 360 nm wavelength (OmniCure S2000). The prepared hydrogels were composed of MuMA alone, MuMA and GelMA (X-Pure® GelMA 160P60 RG, Rousselot, USA), or MuMA and glycerin. In the case of MuMA alone and MuMA-glycerin hydrogels, the MuMA hydrogel precursor and LAP were mixed and dissolved using vortex mixture at RT, and then glycerin was added to the mixture. GelMA hydrogels were prepared by mixing GelMA hydrogel precursor with LAP, vortexing at RT, and then incubating the mixture in the oven at 80 ^0^C for 15-30 minutes until complete dissolution. In the case of GelMA-MuMA hybrid hydrogels, GelMA-LAP, and MuMA-LAP, solutions were prepared separately as described above at the required concentrations and vortexed together just before being cross-linked. The characterized hydrogels were prepared using MuMA alone (Mu), GelMA alone (Ge) or mixture of both (Ge-Mu) at different percentages and ratios indicated by the number preceding the hydrogel abbreviated name (xGe-yMu). In addition, MuMA-glycerin hydrogel was prepared at 20% percentage each and 1:1 ratio (20Mu-20Gly).

### Hydrogel characterization

#### FTIR Spectroscopy

Fourier Transform Infrared (FTIR) measurements were carried out to confirm the cross linking of GelMA and MuMA hydrogels. FTIR spectra were acquired using Attenuated Total Reflectance (ATR) with sample analysis accessory (Nicolet iS5, Thermo Fisher Scientific, USA). Spectra were cumulatively collected for 16 scans using Omnic software by Thermo Fisher Scientific from 400 to 4000 cm^-1^ wavelength. The percent transmittance vs wave number spectra were replotted using OriginPro 2018 software post processing the spectra with baseline correction.

#### SEM imaging and the pore size

The network structures of MuMA, GelMA and their hybrid mixtures were compared using scanning electron microscope (SEM) imaging. The freshly prepared UV-crosslinked hydrogels were freeze-dried, then fixed to the SEM stubs using double-sided conductive carbon tape. Iridium with a thickness of 22 A° was sputter-coated on the surface of the samples in a high vacuum using a sputter deposition and etching system (South Bay Technology, USA). Supra 40 VP scanning electron microscope (Zeiss, Germany) was used to capture the images at an accelerating voltage of 14 kV. The pore area was estimated using Image J software (Image J 1.53K, NIH, USA). The average pore area was calculated from pores with clear boundaries and different representative sizes (n=25).

#### Swelling ratio

The swelling behavior of the hydrogels was evaluated by placing pre-weighed (W0, initial weight) freeze-dried hydrogel discs in a 12-well plate filled with 3 mL DiDW. The plates were wrapped with parafilm and incubated at 37 °C. For sampling; the hydrogel discs were removed from the buffer solution, the excess liquid was blotted gently using paper tissue, and the hydrogel weight was registered (Wt). Hydrogel discs were then re-incubated in fresh DiDW. The results are presented as the swelling ratio of the tested hydrogels versus time, where the swelling ratio at time t is given by Wt/W0. Samples were collected at predefined time intervals: 2, 5, 10, 30, and 60 mins, followed by sampling at 2, 3, 6, 10, and 24 hours for MuMA and at 1, 3, 6, 10, and 24 hours for hybrid hydrogels.

#### Degradation rate

The degradation behavior of the hydrogels was evaluated by placing pre-weighed (W0, initial weight) freeze-dried hydrogel discs in a 12-well plate filled with 3 ml of 10 μg/mL collagenase enzyme solution (Collagenase Powder, Type II, Gibco™, USA). The plates were wrapped with parafilm and incubated at 37 °C. At predefined time intervals (1, 3, and 6 days for MuMA hydrogels and up to 13 days for the hybrid hydrogels), the enzyme solution was pipetted out, and the hydrogel was washed 3 times with DiDW. The excess of the washing liquid was gently removed by blotting, and the 12 well plates containing the hydrogel discs were freeze-dried, and their weight was registered (Wt). The hydrogel discs were then incubated with a fresh enzyme solution for the next data point. The results were presented as the percent remaining of the tested hydrogels where the percent remaining at time t = ((W0-Wt)/W0) x100.

#### Gelation kinetics and rheological characterization

The gelation kinetics were determined by simultaneous measurement of the elastic (G’) and the viscous (G’) moduli under UV exposure for crosslinking. Gelation time sweep was obtained using a rheometer with an 8 mm upper plate diameter (Anton Paar, MCR 302) and UV radiation source of 320-500 nm wavelength (OmniCure, S2000, Excelitas Technologies, USA). For this purpose, 10% (w/v) MuMA hydrogel precursor and 1% (w/v) LAP were dissolved in PBS, then 80 μL of the mixture was placed on the lower plate of the rheometer and covered with the rheometer cap to minimize solvent evaporation. The time sweep test for G’ and G’ was carried out at 10% strain and 1-Hz frequency at 25 ^0^C. During the optimization of the gelation conditions, it was observed that a minimum power of 10 mW/cm^2^ is required for the complete gelation of MuMA alone. The isothermal time sweep of G’ and G’ was conducted for about one minute before starting the exposure to the UV light to establish the measurement baseline.

For the rheological characterization; a rheometer plate-plate configuration with an 8-mm upper plate diameter was used for all the measurements. Hydrogel discs were prepared by pipetting 100 μL of the hydrogel precursor mixture into a PDMS mold measuring 8 mm in diameter and 2.8 mm in depth. The hydrogel was obtained by exposure to UV radiation of different energy and for different exposure periods specified in the results section. The storage modulus was measured as an indicator of the stiffness of the hydrogel. The frequency sweep was carried out using frequency values ranging from 0.1 rad/s to 100 rad/s and at 1% strain (within the linear viscoelastic range, LVER, as determined from the amplitude sweep) at either 25 or 37 ^0^C. The results were presented as G’ and G’ versus the frequency. The temperature sweep was carried out between zero and 50 ^0^C at 2 degrees/minute heating rate, 1% strain, and 1 Hz frequency. The relaxation modulus at 10% strain and 25 ^0^C was also measured. In addition, the average phase shift (loss factor) was calculated for GelMA, MuMA, and GelMA-MuMA hydrogels using Eq. 1.

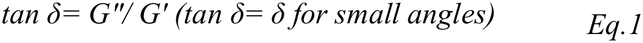

#### Thermal analysis

Differential scanning calorimetry (DSC) was performed with a DSC 823e (Mettler, Switzerland) using aluminum pans that were both pierced and hermetically sealed, each containing 5 mg of the dried hydrogel sample.

Analyses were performed over temperatures ranging from -60 to 400 °C with a heating rate of 10 °C/min, under dry nitrogen purge at 80 mL/min. Identical empty pans were used as the reference. DSC thermograms were obtained for 20Mu, 20Mu-20Gly, 7Ge-7Mu, 7Ge-10Mu, 10Mu, and glycerin alone. To understand the presence of multiple peaks, the samples were heated over a temperature range of 25 to 150 °C, cooled down to ambient temperature, and then reheated over a temperature range of 0 to 400 °C. Thermal gravimetric analysis (TGA) was carried out using TGA (Mettler, Switzerland) with the sample placed in sealed ceramic pans. Analyses were performed over temperatures ranging from -60 to 400 °C with a heating rate of 10 °C/min, under dry nitrogen purge at 80 mL/min. TGA analysis was carried out for 20Mu, 20Mu-20Gly, and glycerin alone.

#### Drug release

The relation between the hydrogel composition and the release profile of the drug was tested using vancomycin hydrochloride (Sigma-Aldrich) as a model drug. Vancomycin is a glycopeptide antibiotic with a molecular weight of 1485 g/mol and water solubility of 100 mg/mL. Drug release from crosslinked loaded films (2 mg vancomycin/film) was measured for 7Ge and 7Ge-7Mu films which were crosslinked under the same conditions. To avoid any dose variations that might be caused by unequal distribution of the drug in the hydrogel precursor mixture; a unit dose of drug-loaded films was used where each film was separately prepared and loaded with the exact dose. For this purpose, a stock solution containing 10 mg/ml vancomycin and 1% LAP in PBS was prepared. From this solution, 200 uL was added to a pre-weighed hydrogel precursor. In the case of GelMA-containing films, the mixture was incubated at 80 °C to produce one phase system before being crosslinked. In the case of hybrid hydrogels, the GelMA-drug mixture was added to the required amount of the solid MuMA and vortexed at RT. The total volume of loaded hydrogel precursor(s) was then transferred using a pipette to a 14 mm diameter PDMS mold and crosslinked at an energy level of 21 mW/cm^2^ for one minute. Four loaded samples were prepared for each experiment. Unloaded films were prepared following the same method, and it was sampled at the same time intervals as the loaded films and served as a blank for each time point. The cross-linked hydrogel films were transferred to a 12-well plate, and 3 ml of PBS were added to each well. The plate was wrapped with a paraffin film to prevent evaporation and placed in a shaker incubator at 37 °C. Samples were withdrawn at intervals extending from 0.3 to 36 hours and replaced by the same amount of PBS that was kept at the same temperature. UV-absorbance measurement at λ= 280 nm was carried out using a UV-spectrophotometer (DS-11, DeNOVIX, USA), and the drug content was calculated using a pre-constructed calibration curve. The stability of the drug upon exposure to heat (80 °C) and UV radiation during the crosslinking procedure was validated using calibrated drug solutions, where no changes in the UV absorbance were observed.

#### Cell Culture and encapsulation

##### Hydrogel biocompatibility test in fibroblast cells, cell viability and metabolic activity assays

BJ (CRL-2522, ATCC) cells were cultured using Dulbecco’s Modified Eagle Medium (DMEM) (Gibco, Inc) supplemented with 10% (v/v) Fetal Bovine Serum (FBS) (Life Technologies), 1% (v/v) penicillin-streptomycin at 37 °C, and 5% CO_2_. Before adding the hydrogels, the cells were cultured in 12-well cell culture plates (Corning Inc, USA) overnight at a resulting density of 1×10^6^ cells/ml. On day 0, the hydrogels, 7Mu, 3.5Ge-3.5Mu, 7Ge-7Mu, and 7Ge suspended in cell culture media were loaded into permeable cell culture inserts of 3 μm pore size (Stellar Scientific, USA) and the inserts were transferred into the well plates containing the cells adhered in the bottom.

Cell viability and cell metabolic activities were assessed on day 2 and day 4 independently using samples from different batches that were similarly treated.To assess viability, the inserts containing the hydrogels were removed and the cells were washed in PBS followed by staining with a LIVE/DEAD viability/cytotoxicity kit (Life Technologies). Briefly, 2 μL/mL ethidium homodimer-1 and 0.5 μL/mL calcein-AM (diluted in PBS) were added to the cell-laden hydrogels, then incubated for 45 minutes at RT in the dark. The cells were washed three times with 1X PBS before imaging. Fluorescence images were collected in both green (live) and red (dead) channels using a Keyence BZ-X810 fluorescence microscope (Keyence Corporation, USA) equipped with a 4X objective. Three randomly chosen areas of 3623 μm x 2718 μm dimensions from each well were imaged and three different wells were imaged per sample. The images were analyzed using FIJI program where cells were manually counted for the number of live and dead cells. The percentage of live cells over the total number of cells (live and dead) were determined for each sample and the control. The average and standard deviation obtained from the three different wells were reported and representative images (cropped and contrast adjusted) are shown in the main text.

Cell metabolic activity was assayed using PrestoBlue™ Cell Viability Reagent (A13261, ThermoFisher). After the inserts containing hydrogels were removed from the wells, media was removed, and the cells were incubated in a mixture of media and PrestoBlue with a 9:1 ratio for 30 min. Media from each condition were pipetted into a 96-well plate for fluorescence analysis using an Infinite M1000 plate reader (Tecan, Switzerland). Average fluorescence intensity from three independent wells per condition were determined reported along with standard deviation.

##### Liver cell encapsulation, viability and metabolic activity assays

HepG2 cells were cultured using Dulbecco’s Modified Eagle Medium (DMEM) (Gibco, Inc) supplemented with 10% (v/v) Fetal Bovine Serum (FBS) (Life Technologies), 1% (v/v) penicillin-streptomycin at 37 °C, and 5% CO_2_. HepG2 cells were added to 3.5Ge-3.5Mu, 7Ge-7Mu, and 7Ge alone in media with 1% LAP at a 2×10^6^ cells/ml concentration. A volume of 100 μL of cell-gel liquid suspension was pipetted into each well in a 96-well plate. The cells were encapsulated in the hydrogels via UV crosslinking for one minute using 21 mW/cm^2^ radiation energy. The 96-well plates were incubated with cell culture media overnight before any experiments were performed. To assess viability, cells were washed in PBS and then stained with a LIVE/DEAD viability/cytotoxicity kit (Life Technologies). Briefly, 2 μL/mL ethidium homodimer-1 and 0.5 μL/mL calcein-AM (diluted in PBS) were added to the cell-laden hydrogels, then incubated for 45 minutes at RT in the dark. The cell-laden hydrogels were washed three times with 1X PBS and then imaged using Zeiss confocal microscope with 5X and 10X macro-objectives on day 1 and day 3. Acquired Z-stack images were Z-projected, and the fluorescent intensity of live cells was measured using Image J software. Cell viability data obtained on day 1 and day 3 were normalized with respect to the day 0 cell number, and the fold increase was calculated for both day 1 and day 3. Cell metabolic activity was assayed using PrestoBlue™ Cell Viability Reagent (A13261, ThermoFisher). Cells were incubated in a mixture of media and PrestoBlue™ with a 9:1 ratio for one hour. Media from each condition were pipetted into a 96-well plate for fluorescence analysis using a plate reader (VARIOSKAN LUX, ThermoScientific).

## RESULTS AND DISCUSSION

### MuMA synthesis and structural characterization

MuMA was synthesized by using a simple, straight forward protocol for methacrylation. Figure 1 shows the sequence of steps involved in preparing the MuMA hydrogel precursor. The PGM used in this work was a very fine dusty powder of low solubility in water at the physiological pH (Barnabás Áron Szilágyi 2020). Consequently, PGM was methacrylated as colloidal dispersion. After methacrylation, mucin was separated from the insoluble components by centrifugation and filtration. Upon lyophilization, the MuMA was produced as a whitish fluffy mass (**Fig. 1**) that dissolves readily in phosphate buffer saline (PBS) at room temperature (RT), forming a clear solution. The solubility of freeze-dried MuMA compared to pure mucin can be attributed to the absence of the insoluble components removed during the centrifugation and filtration steps. The hydrogel precursor prepared following the protocol described above ensured that only the soluble part of mucin was used in the hydrogel preparation. The degree of methacrylation was then determined using NMR analysis as described in the next section. In addition, the presence of methacrylic acid was examined using the FTIR spectral analysis.

**Fig. 1:**
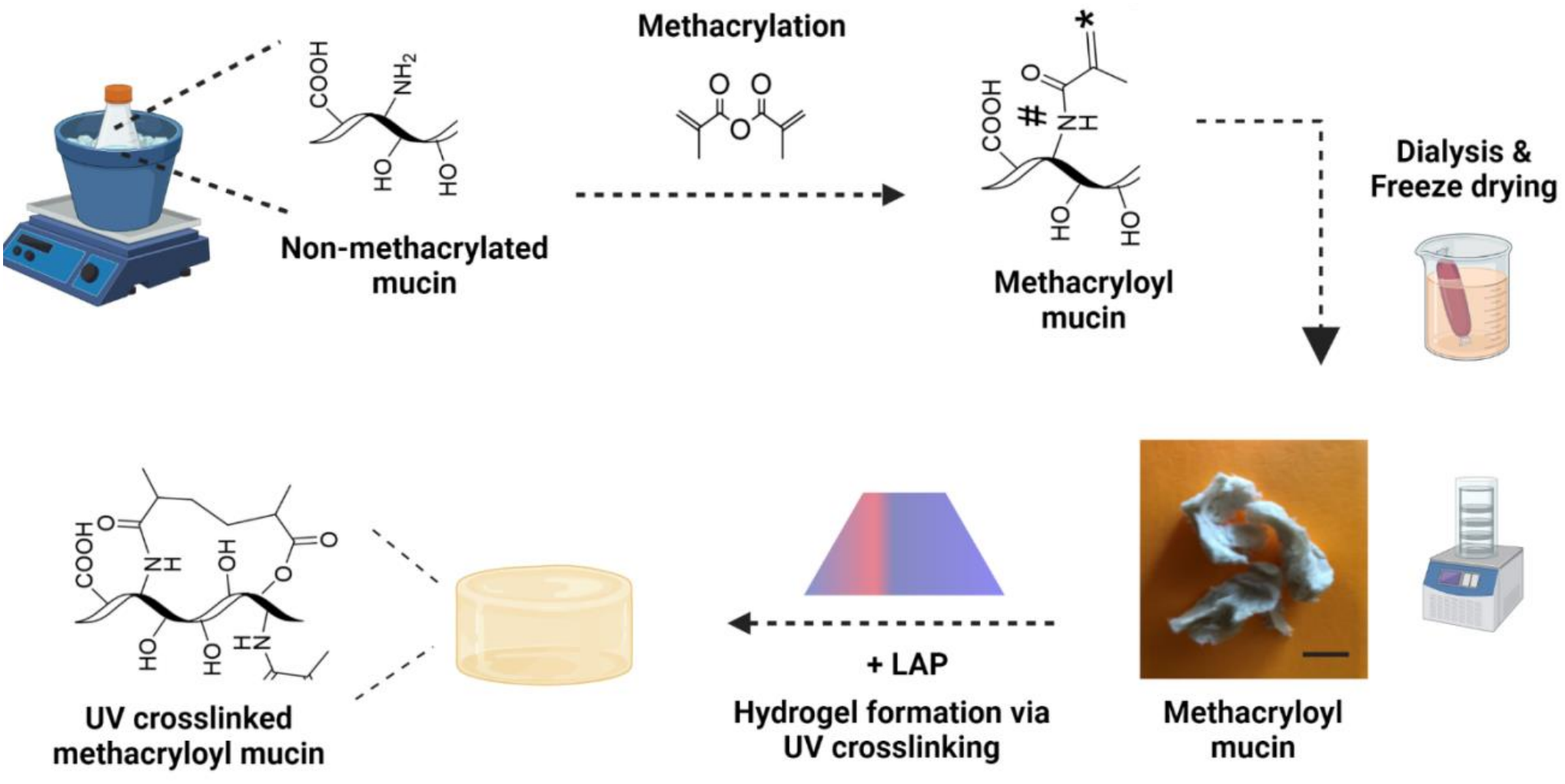
The main steps of mucin methacryloyl process and the subsequent hydrogel formation using UV crosslinking.

#### NMR characterization

Figure 2A describes the suggested mechanism of the mucin methacrylation process. Mucin is a heterogeneous glycoprotein (Carraway 1991) containing lysine as a basic amino acid that offers amine moieties and sugars that offer hydroxyl moieties (Liu 2021). Crosslinking can occur either at the core protein or brush oligosaccharides (Kun Jiang 2021). Serafim *et al*. (Serafim 2018) suggested that the amino group of lysine is more prone to reacting with acids than hydroxyl groups. This might be due to the relative stability of intramolecular hydrogen bonding among sugars’ hydroxyl group; the terminal amino group of lysine is ionized at the physiological pH and serves as a nucleophile at pH 8.0 in the reaction vessel (Liu 2019). The pKa of the terminal amine of lysine is around 9.0 (Pahari 2019) so at the pH of the reaction vessel, around 10% of the amino groups are in the basic form. On the other hand, the ammonium groups cannot act as a nucleophile, but they donate basic amines when the latter are consumed in the reaction (Takats 2001). Thus, the lysine terminal amine plays the role of a modification site with a suitable acylating agent. Methacrylic anhydride was used here due to its ability to remove a good leaving group (Hu 2005). The reaction releases methacrylic acid, which can reduce the pH. However, the reduction of the pH increases the ionization of the terminal amines and consequently suppresses the reaction process. Thus, continuous adjustment of the reaction pH with an alkaline solution is performed to maintain the desired pH throughout the methacrylation process (Fakhari 2013, Hong 2013).

**Fig. 2.**
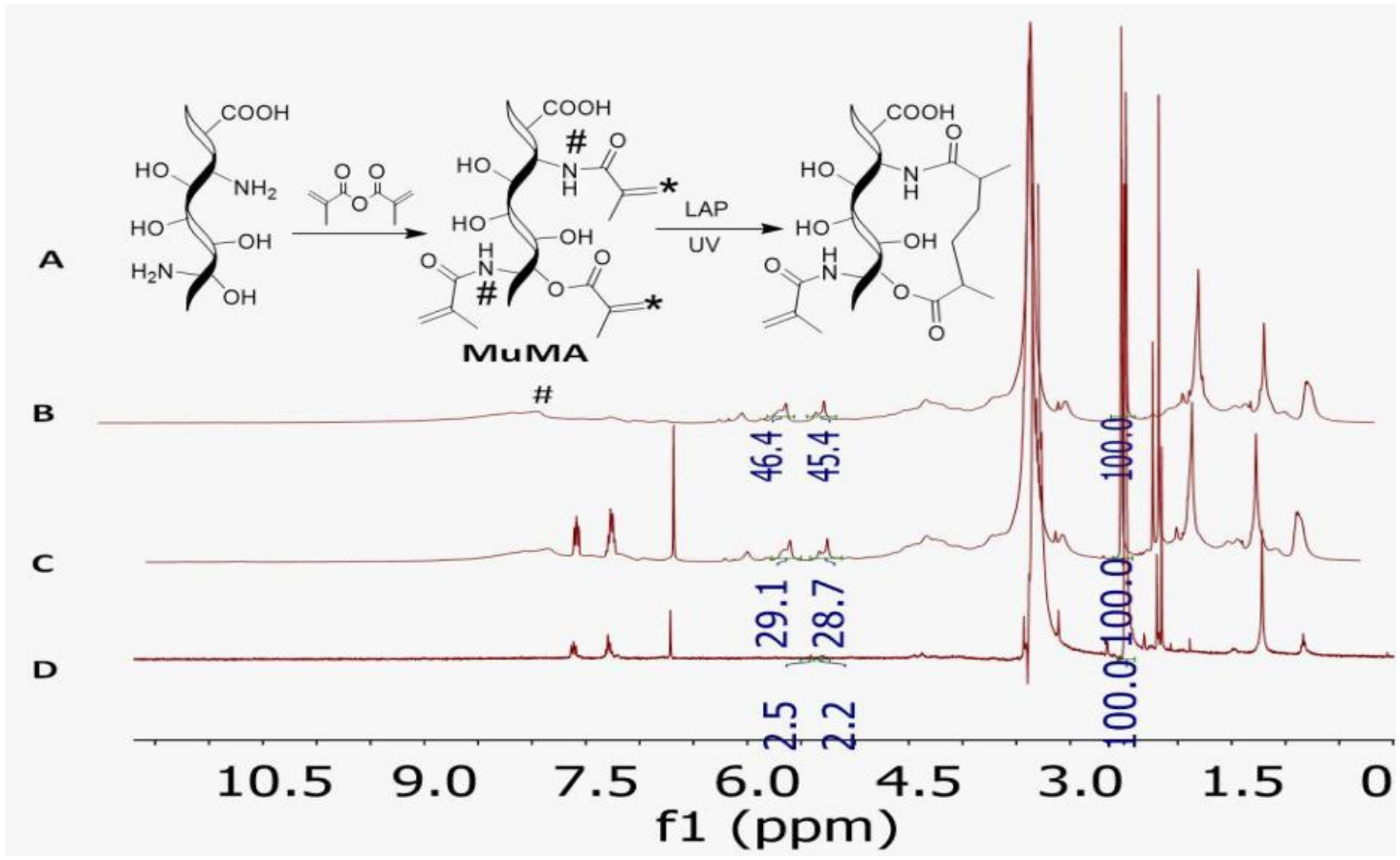
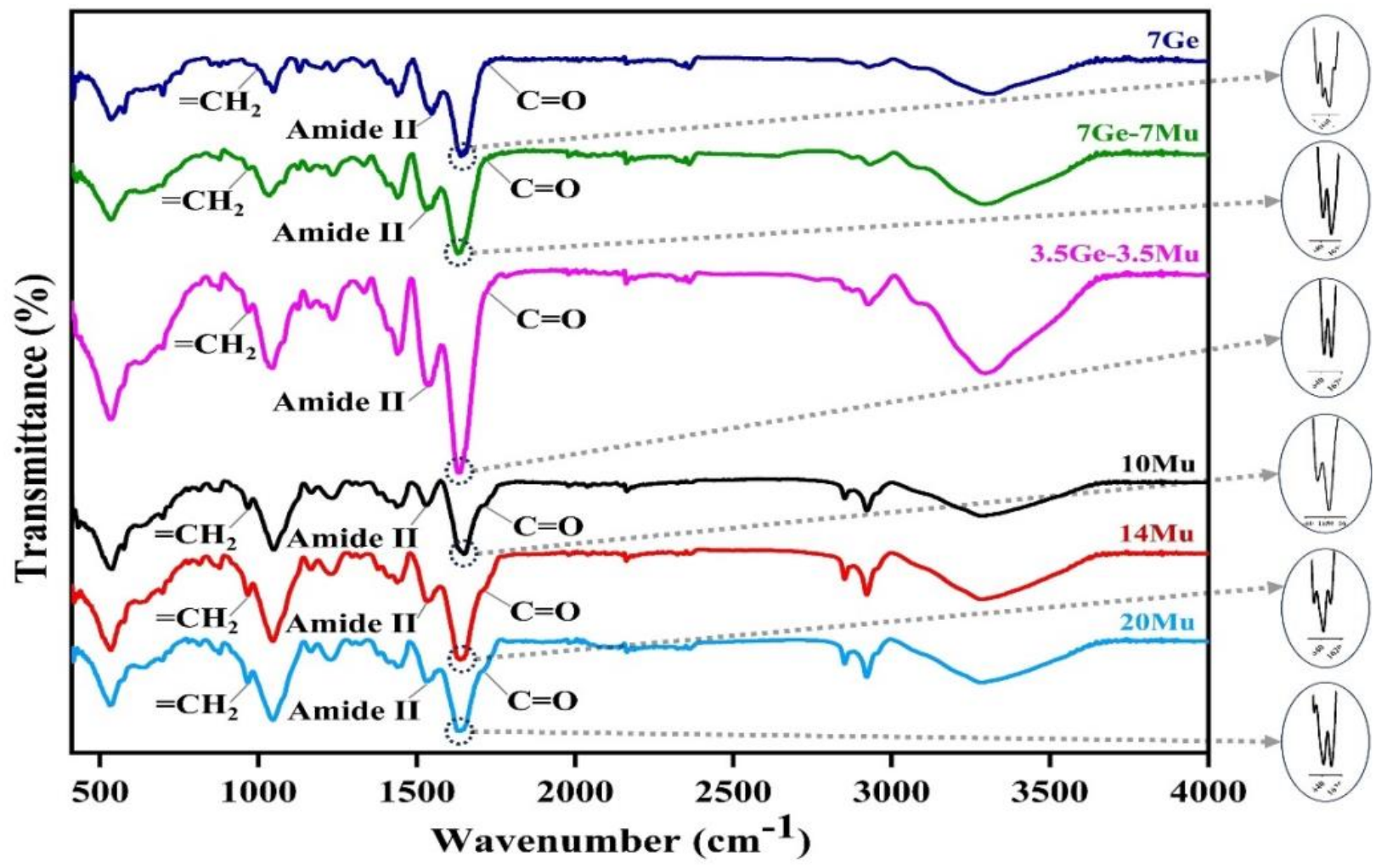
Upper panel: NMR spectra for mucin methacryloyl. **A:** Methacrylation (#) of lysine moieties in mucin with methacrylic anhydride. **B**: NMR of physical mixture of MuMA and the photoinitiator (LAP) without crosslinking. **C:** NMR of crosslinked MuMA. Double-bond hydrogens almost disappeared. **D:** NMR spectrum of MuMA in DMSO. The appearance of double bond hydrogens (*) at around 5.5 ppm, an acrylic acid methyl group at 1.8 ppm, and hydrogens of amide linkage at around 7.0 ppm are proof of mucin modification in MuMA. **Lower panel:** FTIR spectra showing signature peaks of 7Ge, 7Ge-7Mu, 3.5Ge-3.5Mu, 10Mu, 14Mu and 20Mu. Zoomed regions show distanced combined peaks (see the text).

Additionally, the pH adjustment with an alkaline solution increases the solubility of liberated methacrylic acid and enhances the reaction to proceed forward according to the mass action principle. It is noteworthy to mention that sugars’ hydroxyl group might react to a certain extent with methacrylic anhydride, but the product is ester rather than amide. Generally, amides are more stable than esters and more thermodynamically favored products (Deslongchamps 1975). As depicted in Figure 2D, the methacrylation was confirmed by observing hydrogens of amide groups resulting from the reaction of lysine with methacrylic anhydride at around 7.0-8.0 ppm (Gellman 1991, Harding 1991). Interestingly, the amide of peptides bonds (Fig. 2A) was not clearly seen in the NMR of mucin because of their heterogeneity in terms of electron properties (**Fig. 2B**); therefore, the signal of amide hydrogens broadened significantly and disappeared (**Fig. 2C**). Additionally, the solubility of mucin was relatively low in aqueous medium and DMSO. At the same time, MuMA had better solubility in both solvents. This might also participate in the disappearance of the peptide hydrogen signals in mucin. The other proofs of methacrylation were the methyl chemical shift at around 1.8 ppm and the double bond hydrogens in the double bond region at 5.0-6.0 ppm (**Fig. 2D**).

The integration of double-bond hydrogens of acrylic acid relative to the DMSO signal in MuMA reduced significantly after cross-linkage with UV radiation. The methyl hydrogens of methacrylic acid were shifted downfield (**Fig. 2C**). According to Equation 3 (Eq.3), the degree of cross-linkage was 96%. Interestingly, the sharp signals of aromatic hydrogens at around 7.0 ppm are due to the hydrogens of lithium phenyl-2,4,6-trimethylbenzoylphosphinate (LAP).

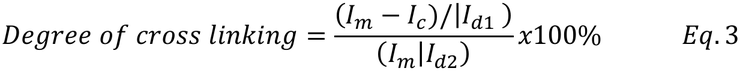

Where *I_m_* is the integration of double-bond hydrogens in MuMA, *I_c_* is the integration of double-bond hydrogens in cross-linked MuMA, and *I_d_* is the integration of the DMSO signal at 2.5 ppm as a reference signal.

As the free radical polymerization was achieved using UV radiation and LAP as the photoinitiator, the reaction proceeded to link methacrylic acid moieties via the conversion of π bonds to more stable δ bonds. This process leads to the polymerization of methacrylic acid molecules. However, the polymerization process not necessarily links only mucin molecules with each other’s. Instead, polymerization might occur within the same mucin molecule. The physical mixture of MuMA and LAP showed intensities for double bond hydrogens exceeding those in the crosslinked polymer by 10-fold (**Fig. 2B and C**), thus demonstrating the effectiveness of the photoreaction in crosslinking methacrylic groups.

### Hydrogel characterization

MuMA hydrogels at different MuMA concentrations and MuMA-GelMA hydrogels at different concentration rations were prepared and characterized with regards to their FTIR spectroscopy, network structure, viscoelastic properties, swelling and degradation behavior. The lowest MuMA concentration required to produce a hydrogel was 5%, but the hydrogels examined in this work were prepared at higher MuMA concentrations of 7%, 10%, 14%, and 20% (w/v), denoted as 7Mu, 10Mu, 14Mu, and 20Mu, respectively, and crosslinked using energy 21 mW/cm^2^ for one and five minutes (KangJu Lee 2020). For 7Mu, one minute of exposure was insufficient to form a hydrogel. MuMA hydrogels were opaque with a faint yellowish color (**Fig. 3A**).

**Fig. 3:**
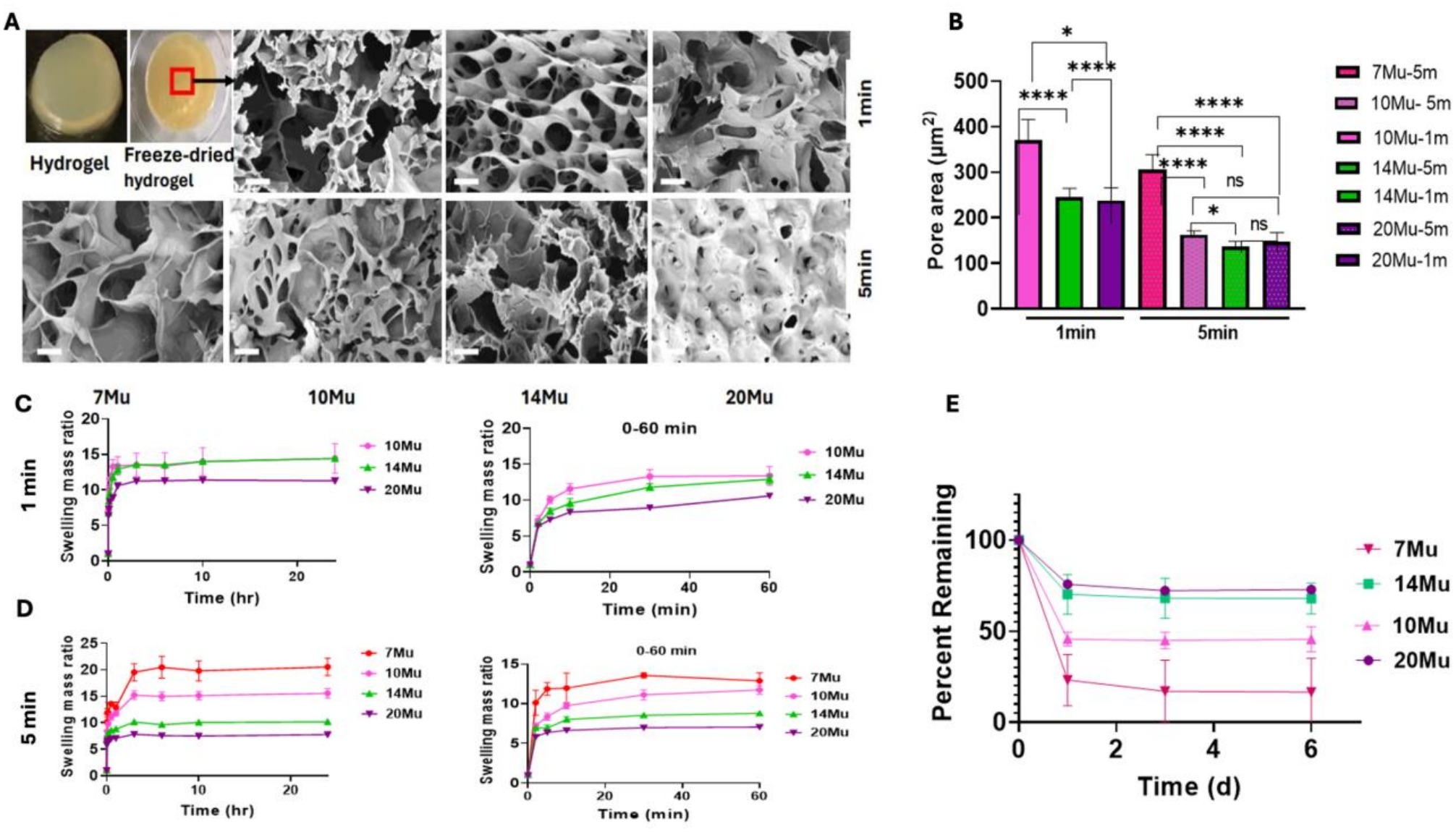
Characterization of MuMa hydrogels prepared at different MuMA concentrations and crosslinked for one or five minutes using 21 mW/cm^2^ energy. **A:** Hydrogel morphology and SEM images (scale= 20 μm). **B:** Hydrogels’ pore sizes obtained from the SEM images. **C** and **D:** Swelling ratio (left: over 24 hrs, right: the first hr.) (Average ± SE, n=2-5) of hydrogels crosslinked for one and five minutes, respectively, and incubated at 37 ^°^C in DiDW. **E:** Degradation profile (Average ± SE, n=3-4) of hydrogels cross linked for five minutes incubated in collagenase at 37 ^0^C. Statistically significant differences were identified using the Student’s t-test. P-values were lower than 0.05 (*), 0.01 (**), and 0.0001 (***) or higher than or equal to 0.05 (not significant).

#### *FTIR* Spectroscopy

FTIR spectroscopy was performed for MuMA hydrogels at different concentrations and for GelMA-MuMA hybrid hydrogels, to examine the interaction between the two polymers. Figure 2, lower panel illustrates that all the examined hydrogels showed the signature functional groups peaks of Mucin, Gelatin and Methacryloyl in the FTIR spectra. The O-H and N-H stretching vibration overlapped peak was observed as a broad peak between 3100–3700 cm^-1^. Peaks at around 2851 cm^-1^ and 2921 cm^-1^ depicted N-H stretching for amine group and C-H stretching representing alkanes respectively. Significant peak around 1630-1660 cm^-1^ on deconvolution (showed in circle besides respective spectra) further distanced the combined peaks; from which peak at around 1648-1654 cm^-1^ refer to Amide I. The extended shoulder at 1708-1721 cm^-1^ from the same peak signifies the carbonyl group (C=O) formed because of methacrylic anhydride modification. Furthermore, the peak representing Amide II at around 1534 cm^-1^ which was slightly shifted to that of the signature peak observed in Mucin at 1554 cm^-1^ (Olăreț, Bălănucă et al. 2021). Peaks at 967 cm^-1^ and 1638 cm^-1^ refer to =CH_2_ deformation and -C=C-stretching vibration respectively and they are key bands for the Methacrylate grafting (Balanuca, Lungu et al. 2014) on Mucin and Gelatin. Moreover, MuMA and GelMA specific C-H deformation vibration was noted at 1378 cm^-1^ and 1438 cm^-1^ (Serafim, Cecoltan et al. 2020). Thus, ATR-FTIR spectroscopy conducted for all the hydrogels of different polymeric combinations and concentrations had signature peaks directing to the presence of Methacrylate in both GelMA and MuMA.

#### SEM imaging and the pore size

The network patterns of MuMA hydrogels, as captured by SEM images are illustrated in figure 3A. The pore size values were slightly heterogenous within the same hydrogel, but there was a trend of increase in the pore size with decreasing both the hydrogel concentration and the irradiation period. The pore areas for hydrogels above 14% (w/v) mucin concentrations showed no significant differences, irrespective of UV exposure time. When the hydrogels were crosslinked for five minutes, the pore area of hydrogels with high MuMA content (10Mu, 14Mu, and 20Mu) was almost similar (average 150 μm^2^) but significantly different (P<0.001) from 7Mu (300 μm^2^) (**Fig. 3B**). Similarly, the pore areas for 14Mu, and 20Mu hydrogels cross-linked for one minute were almost similar (average 250 μm^2^) but significantly smaller (P<0. 01) than the pore size of 10Mu hydrogel (380 μm^2^).

#### Swelling ratio

The swelling mass ratio of MuMA hydrogels, calculated as the total hydrogel weight at different times divided by the initial weight of the freeze-dried hydrogel, is shown in figure 3C and 3D. Hydrogels became less opaque as the MuMA content decreased or upon excessive hydration, as observed during the swelling experiments. For all tested MuMA hydrogels, equilibrium swelling was achieved within the first three hours of incubation irrespective of the polymer concentration or the irradiation period. Hydrogels irradiated for five minutes (**Fig. 3D**) illustrated that the swelling ratio decreases with the increase in the MuMA content. This trend was less obvious in hydrogels irradiated for one minute (**Fig. 3C**) where 20Mu showed lower equilibrium swelling ratio compared to 14Mu and 10Mu which exhibited almost similar swelling equilibrium values. The decrease in the swelling ratio of MuMA hydrogels with increasing MuMA content can be explained by the formation of a dense cross-linked network structure. This finding is important in the design of MuMA based medicated carriers with tailored swelling and consequently, drug release behavior.

#### Degradation rate

The enzymatic degradation using a collagenase enzyme was conducted to validate the suitability of MuMA hydrogels for *in vivo* applications. The hydrogels’ degradation rate was 7Mu > 10Mu > 14Mu and 20Mu (**Fig. 3E**). The degradation rate is fast for hydrogels of lower solid content because increasing MuMA concentration results in a denser network with a lower surface area exposed to enzymatic interaction.

#### Gelation kinetics and rheological characterization

The gelation kinetics profile (**Supplementary Figure 1**) shows that the plateau in the elastic modulus was reached after about 10 minutes of UV exposure, indicating the completion of the gelation process. The viscoelastic properties of the hydrogels were measured using a parallel plate rheometer. The storage and loss moduli, relaxation modulus, and the effect of temperature on the viscoelastic behavior were investigated. The frequency sweep of the hydrogels (**Fig. 4A-F**) showed an increase in G,’ and G’ values with increasing MuMA concentration. Significant differences in both G’ and G’ values were observed in hydrogels exposed for five minutes to UV radiation compared to one-minute exposure where minor differences were observed (**Fig. 4C and F**). Temperature sweep data showed the stability of G’ and G’ values for 10Mu, 14Mu, and 20Mu hydrogels up to 50 °C and a minor fluctuation was observed in the case of 7Mu (**Fig. 4G, H, J, and K)**. On the other hand, the loss modulus (G’) showed a non-significant decrease with the temperature increase in all hydrogels. The relaxation modulus, G(t), profiles (**Fig. 4I and L**) clearly showed the effect of crosslinking period and MuMA concentration on G(t). After one minute of UV exposure, the G(t) value was constant for 14Mu and 20Mu hydrogels, while a decrease in G(t) value was observed for 10Mu hydrogel upon five minutes of UV exposure. However, 7Mu hydrogel showed a decreasing G(t) value over time irrespective of exposure time. Altogether, these results suggest the predominance of elastic behavior in the case of high MuMA concentration and longer crosslinking periods. On the contrary, the G(t) value for 7Mu hydrogel reached almost zero, indicating a very weak elastic network structure.

**Fig. 4:**
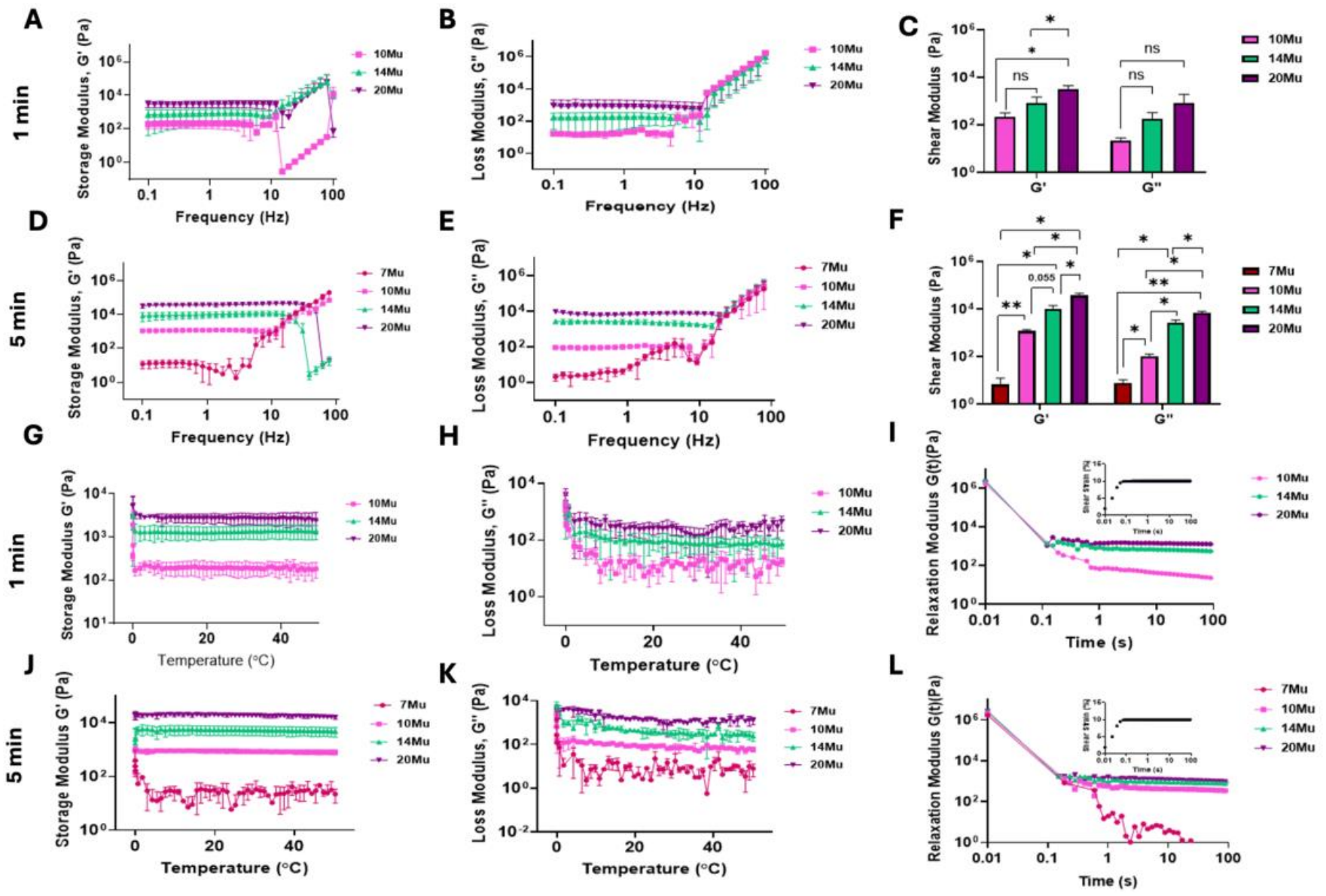
Rheological characterization of MuMA hydrogels at different concentrations, crosslinked using 21 mW/cm^2^ energy for either five minutes (7Mu, 10Mu, 14Mu, and 20Mu) or one minute (10Mu, 14Mu and 20Mu). The measurements were carried out at 25 °C. **A-F:** G’ and G’ values obtained from frequency sweep measurements at 1 % shear strain, (Average ± SE, n=3). **G, H, J** and **K:** G’ and G’ values obtained from temperature sweep measurements at 1% shear strain and at 1 Hz frequency (Average ± SE, n=3). **I** and **L**: The relaxation modulus was measured at 10% shear strain and 1Hz frequency (Average ± SE, n=3). The inserts represent the constant strain applied to the sample. Statistically significant differences were identified using students’ t-test with the denotations indicating p-values lower than 0.05 (*) and 0.01 (**).

Interestingly previous reports suggest that physical crosslinking occurs naturally in hydrogels (Hoffman 2012, Sheila Maiz-Fernández and Nagore Gabilondo 2020), but more particularly in mucin hydrogels (Lynda A. Sellers 1988, Lynn M. Soby 1990, Oliver W. Meldrum and Michael A. McGuckin 2018). In addition to the UV crosslinking, the contribution of physical crosslinking in the high-content MuMA hydrogels is thought to be responsible for thermal stability and the relatively high equilibrium value of the relaxation modulus.

### Role of additives in modifying MuMA hydrogel properties

MuMA hybrid hydrogels containing either a second gel precursor (GelMA) or a plasticizer (glycerin) were also prepared and characterized. The results indicated that GelMA-MuMA hydrogels can be obtained using shorter irradiation period when compared to MuMA hydrogel at the same total concentration. A minimum concentration of 7% MuMA (7Mu) and five minutes of exposure to 21 mW/cm^2^ energy were needed to form a hydrogel of equivalent consistency to that produced after one minute of radiation exposure using 3.5% of each hydrogel precursor (3.5Ge-3.5Mu). The hydrogel formation at this low concentration of the individual polymers might be explained by forming an interpenetrating GelMA-MuMA hydrogel network since both hydrogel precursors were crosslinked simultaneously (J. Aleman 2007). This interaction showed great impact on the hydrogel properties. The extent of the network formation in hydrogels might be described using the phase shift values. MuMA hydrogels showed a higher phase shift value than GelMA hydrogels (**Fig. 5A**), irrespective of the hydrogel formulation parameters, indicating a weaker network which might be related to the presence of the deprotonated carboxylate groups in MuMA hydrogels at a pH of 7.4 (Mimpin Ginting 2019). A significant decrease in the phase shift was observed in the hybrid hydrogels upon the addition of GelMA compared to MuMA alone. Using a mixture of GelMA and MuMA in the hybrid hydrogel seems to increase the networking in the hydrogel and consequently decrease the phase shift values; both polymers most probably formed an interpenetrating polymer network (L. F. Wang 2002, J. Aleman 2007). The swelling behavior of MuMA hydrogels was also modified by the addition of GelMA, the swelling mass ratio at 24 hours for 3.5Ge-3.5Mu and 7Ge-7Mu were 47 and 16.5 (**Fig.5 B**) compared to 20 and 10 for 7Mu and 14Mu respectively (**Fig.3 D)**. In addition, 3.5Ge-3.5Mu showed a significantly large pore size (Supplement Figure 2) of 690 ± 98 µm^2^ compared to 305 ± 33 µm^2^ for 7Mu (**Figure 3A**). The relationship between pore area and porosity and the cell behavior was studied extensively (Nicole J. Darling 2016, Valeria Nele 2020), where it was suggested that a larger pore size is advantageous for cell proliferation. This observation was attributed to the ease of spreading experienced by the cells cultured in the larger pore size matrix, which leads to higher cell proliferation rates, while the smaller pore sizes limit the formation of large cell aggregates and reduce cell differentiation (Choong 2013). Having low concentration of the hydrogel precursor is essential for creating a cell-promoting microenvironment by controlling the pore size, which affects oxygen and nutrient diffusion, migration, and intercellular interactions (Iliyana Pepelanova 2018). Adjustment of the pore size was also reported to contribute to the formulation of hydrogels having a controlled drug release (Li-Fang Wanga 2003, Muhammad Suhail 2021). The effect of additives on the pore size might also partially explains the degradation profile of hybrid hydrogels (**Fig. 5C**). MuMA hybridization with GelMA increased the degradation rate compared to MuMA alone hydrogels (**Fig. 3E and Fig. 5C**). The higher degradation rate of MuMA-GelMA hybrid hydrogels might be related to the larger surface area, in the hybrid hydrogels, that is available for degradation interaction. Adding glycerin into MuMA hydrogel produced a flexible transparent film (Supplement Figure 3) as glycerin interacts with the polymer chains, weakening the interchain interaction and increasing the polymer chains’ mobility, as a direct result of this effect, complete loss of the LVER was observed when the frequency sweep was carried out for 20Mu-20Gly hydrogels at 37 ^0^C. Glycerin also caused a slight increase (about 4%) in the swelling mass ratio of 20Mu-20Gly hydrogel compared to 20Mu alone. This effect was expected because of the hydrophilic nature of glycerin (Guohou Sun 2018).

**Fig. 5:**
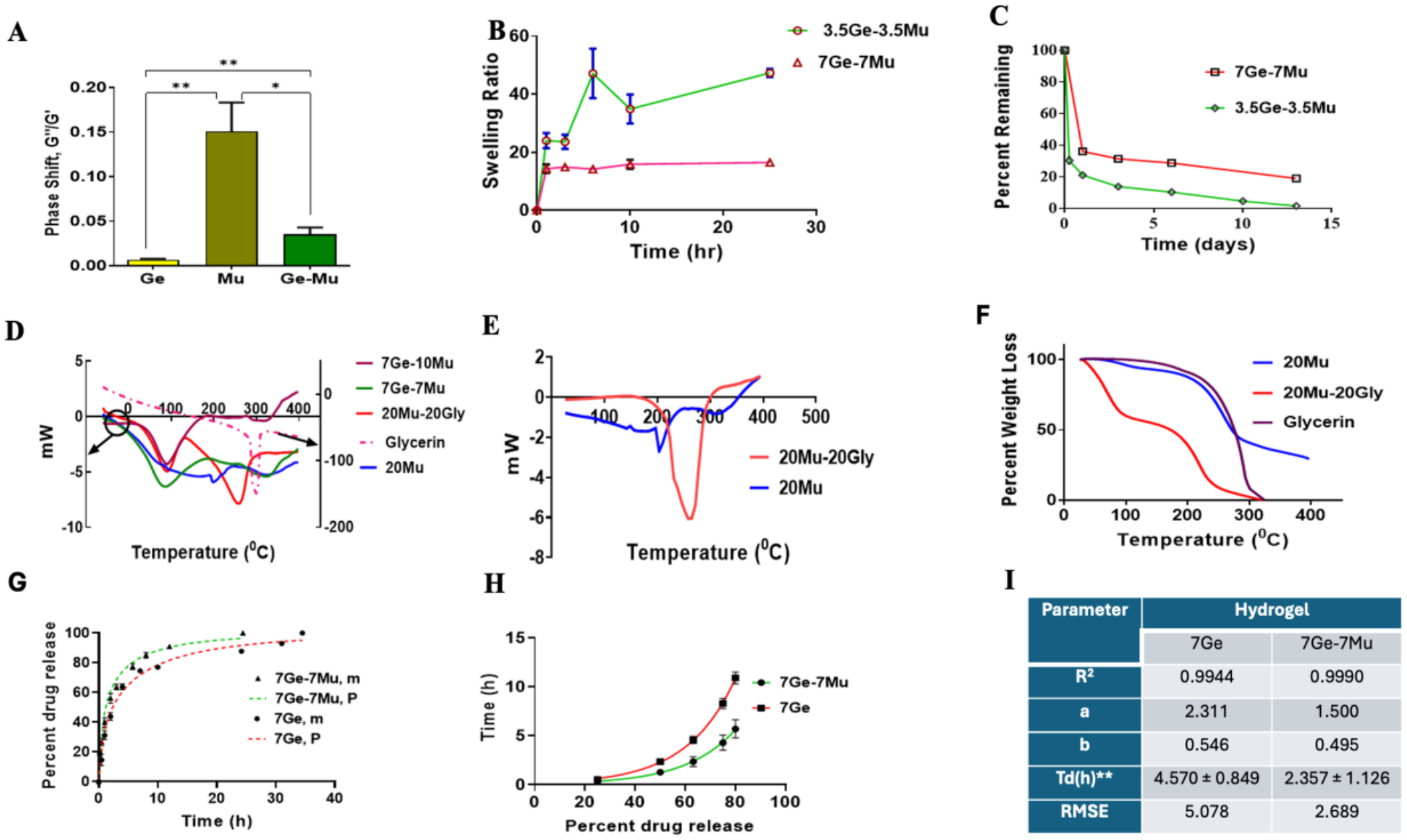
Hybrid hydrogel characterization. **A:** Phase shifts (Average ± SE) of Ge hydrogels (n=3), Mu hydrogels (n=18), and Ge-Mu hydrogels (n=15). Statistically significant differences in the phase shift values were identified using the Welch test. **B:** Swelling ratio (Average ± SE, n=2-5) of 3.5Ge-3.5Mu and 7Ge-7Mu. **C:** Degradation (Average ± SE, n=3-4) of 3.5Ge-3.5Mu, and 7Ge-7Mu hydrogels incubated in collagenase at 37 ^0^C. **D:** DSC thermograms (mv versus temperature) of 20Mu alone, 7Ge-10Mu, 7Ge-7Mu, 20Mu-20Gly, and pure glycerin. **E:** DSC thermograms of heated-cooled-reheated samples of 20Mu and 20Mu-20Gly. **F:** TGA thermograms for 20Mu alone, 20Mu-20Gly, and pure glycerin. **G-I:** Vancomycin release (Average ± SE, n=3-4) in PBS from 7Ge-7Mu, and 7Ge hydrogels measured at 37 ^0^C. **G:** The experimentally measured VM release (n=3) percentage, m vs. time and the corresponding best-fit, P, performed with the use of the Weibull model, Equation (4). **H**: The exponential distributions of VM release time. **I:** Weibull Model Parameters of drug release from 7Ge and 7Ge-7Mu films.

#### Thermal analysis

The DSC thermograms for representative hydrogel formulations are shown in figures 5D, 5E and 5F. The DSC thermogram of 20% Mu MA, depicted in figure 5D, reveals two broad peaks at 169°C and 327°C, accompanied by an endothermic minimum at 201°C. The notable peaks corresponded to a weight reduction of 12 % and 57 % during two intervals at 50 - 200°C and 200 - 400°C, respectively, as depicted in the TGA thermogram of 20% MuMA (**Figure 5F**). The depletion of water from dehydration was associated with the initial stage of weight reduction of 12 %, which occurred between 20 and 200°C. The depolymerisation of MuMA, followed by degradation at elevated temperatures, caused weight loss of 57% between 200 and 400°C (Li-Fang Wang 2003, Muhammad Suhail 2021). The sample continued to decompose until the end of the run at 400°C.

The interaction between MuMA and either GeMA or glycerin was investigated through DSC thermal analysis. The thermogram of pure glycerin exhibited an endothermic peak at approximately 300°C (Figure 5D), aligning with glycerin’s boiling point and concurrent evaporation, as indicated by the weight loss in Figure 5F.

In the analysis of hybrid hydrogels, specifically 7Ge-10Mu and 7Ge-7Mu, distinct transition peaks were identified, with the initial broad endothermic peak occurring at 90°C (Fig. 5D). The endothermic melting peak of 20Mu at 201°C was not discernible in these hybrid hydrogels. A potential explanation for both observations may be the molecular interaction between GeMA and MuMA, which facilitates a reduced binding of water molecules to MuMA, consequently leading to water evaporation at a lower temperature (90°C) (Yoshinori Tamai 1996, Yoshinori Tamai 1996). Likewise, glycerin engages with MuMA, facilitating the liberation of water molecules, which can then evaporate at a reduced temperature. Two endothermic peaks were observed at 94°C and 257°C in the thermogram of 20Mu-20Gly (**Fig. 5D**).

The 20Mu-20Gly TGA thermogram exhibited two distinct phases of weight reduction. Water evaporation resulted in a 44.4% reduction in weight during the initial phase, aligned with the first endothermic peak, which occurred between 25 and 130°C (**Fig. 5D**). The volume of water in this sample was considerably greater than that retained in the 20% MuMA sample. A weight reduction of 56.5% was noted at the second stage, aligning with the second endothermic peak of the DSC thermogram between 150 and 324°C (**Fig. 5D**). This observation demonstrated that glycerin interacts with MuMA in the 20Mu-20Gly sample, leading to simultaneous water evaporation and MuMA depolymerisation at reduced temperatures. Thus, thermal analysis revealed the interaction between GeMA and MuMA, as well as between MuMA and glycerin.

Figure 5E illustrates the DSC thermogram of 20% Mu MA after being heated to 150°C in a perforated crucible, then cooled to ambient temperature, and reheated to 400°C. The thermogram exhibits an endothermic minimum at 141–142°C, indicating the glass transition temperature (Tg), succeeded by an exothermic peak at 149°C, which indicates the crystallization temperature. The presence of Tg in the DSC thermogram signifies the amorphous properties of the dried MuMA sample. The melting of the partially crystalline Mu MA is indicated by an endothermic peak at 203°C.

The 20Mu-20Gly hydrogel was subjected to heating at 150°C in a DSC-perforated crucible, followed by cooling to room temperature and then reheating to 400°C (**Fig. 5E**). Following this heating-cooling-heating cycle, the DSC thermogram of the 20Mu-20Gly mixture exhibited an endothermic peak at 264°C, which is lower than the melting point of pure glycerin at 300°C. This observation indicates a significant molecular interaction between MuMA and glycerin.

Furthermore, the DSC thermogram of the reheated 20Mu-20Gly hydrogel showed that the melting endotherm of 20Mu, which had previously been detected at 203°C, was no longer present and had merged with the glycerol endothermic peak. This alteration suggests a molecular interaction between the two components.

#### Drug release

Hybrid 7Ge-7Mu and 7Ge hydrogels were tested for drug release behavior using vancomycin (VM) as a model drug. The results are presented in figures 5 G-I. The kinetics of drug release were assessed using model dependent techniques such as first-order model, Higuchi model, Korsmeyer-Peppas model, Hixson and Crowell model and the Weibull model. For both hydrogels, Weibull model, equation 4, offered the best correlation coefficients (R^2^>0.99) and lower root mean square error (RMSE) values (2.7 and 5.1 for 7Ge-7Mu and 7Ge respectively). The parameters associated with this model **(Eq. 4)** are parameter (a), which specifies the process time scale; localization parameter (Ti), which denotes the release process lag time; and parameter (b), which describes the dissolution curve shape. Equation 4 shows that the curve’s shape for b = 1 is precisely the same as the shape of an exponential profile with constant k = 1/a.

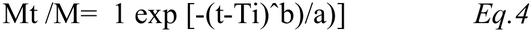

Where Mt is the amount of drug being released as a function of time t. M is the total amount of drug being released. A curve with a value of “b” greater than one has a sigmoidal shape with a turning point; in contrast, a curve with a value of “b” less than one would have a steeper increase than one when “b” equals 1. The parameter “a” can be replaced by the more informative release time, “Td” parameter (**Fig. 5I: table**). The “Td” parameter represents the time interval necessary to release 63.2% of the drug present in the film (Justyna Kobryń 2017).

Figure 5H shows the suitability of equation 4 in the prediction of the extent of drug release over the entire experimental range for both hydrogels. The steep exponential release characteristics of both films, with “b” values smaller than 1 (0.495 and 0.546 for 7Ge-7Mu and 7Ge respectively) indicating Fickian diffusion in both hydrogels. The “b” value was related to the extent of the disorder in the hydrogel structure. Hydrogels with lower disorder showed higher “b” values (Vasiliki Papadopoulou 2006). The two hydrogels showed a slight difference in their “b” values which is in agreement with the difference in the ratio of the viscous to elastic moduli (phase shift, **Fig.5 A**).

Large differences were observed in the scale parameters “a” (2.311 and 1.5 for 7Ge and 7Ge-7Mu respectively) and the parameter “Td” (4.570 ± 0.849 and 2.357 ± 1.126 for 7Ge and 7Ge-7Mu respectively, P < 0.10. Mean ± SD). This difference agrees with the 7Ge hydrogel slower release rate compared to 7Ge-7Mu hydrogel. Further confirmation is also shown in figure 5 H that describes the exponential distributions (R2>0.995) of the drug release patterns.

Drug release kinetics are governed mainly by the dissolution or degradation of the hydrogel matrix. The Fickian diffusion of the drug through the hydrogel network and hydrogel swelling is expected to slow down the drug release processes (Sarita R. Shah 2014). As a hydrophilic drug, VM is expected to be trapped in the liquid-filled pores of the network formed in the swollen hydrogel. Vancomycin was reported to interact via its positive charge with mucin sialic residue causing mucin aggregation (Vlad Dinu 2020).

This type of interaction suggests a delayed release of VM from mucin containing hydrogels, a case that was not observed in this work. Similarly, vancomycin loading into negatively charged biodegradable polymeric microparticles was found lower than expected based at its positive charge (Sarita R. Shah 2014). It is worth mentioning here that VM has multitude of pKa values, with a net charge of only +0.7 at pH 7.4 (Vijan 2009), which might explain the unexpected weak interaction with mucin.

#### Cell Biocompatibility Assay

Biocompatibility of the GelMA-MuMA hybrid hydrogel was tested using fibroblast cells. Four different hydrogels were tested: GelMA-MuMA hydrogels at two compositions, GelMA alone and MuMA alone: 7Mu, 3.5Ge-3.5Mu, 7Ge-7Mu, and 7Ge. None of the hydrogels impacted cell viability, thus confirming the high potential of GelMA-MuMA for tissue engineering applications (**Fig. 6**). The cell viability was assessed on day 1 and day 4 using Live/Dead assay as shown on figure 6A. All samples show 85 to 95 percentage viable cells for both day 2 and day 4 (**Fig. 6 A-B**). For better understanding; cell metabolic activities on day 1 and day 4 were performed independently. After four days of culture, a significant increase in cell metabolic activity was observed for all 4 samples (p > 0.05) (**Fig. 6 C**).

**Fig. 6:**
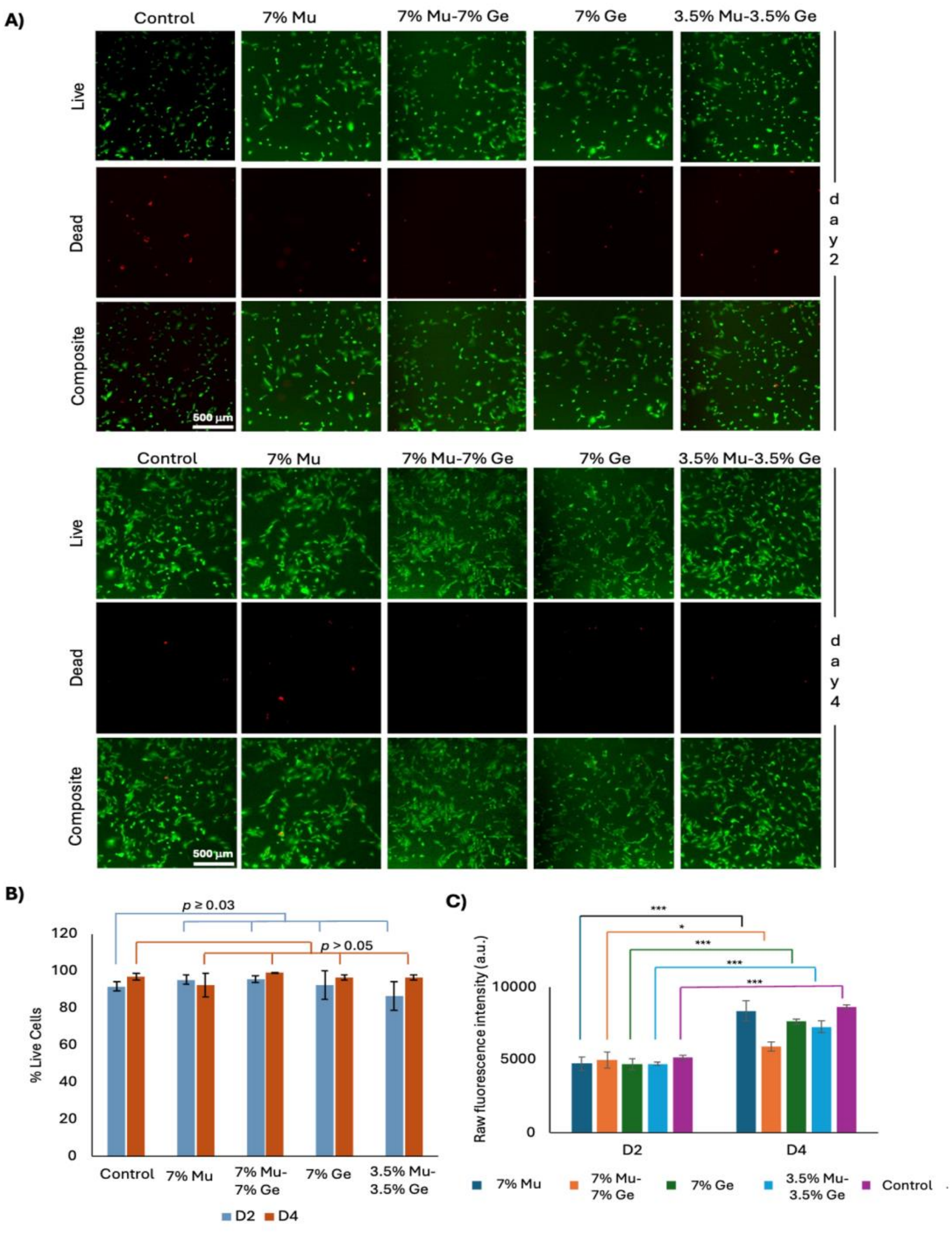
Biocompatibility assay performed on day2 and day 4: (A) Live (green) and dead (red) images of BJ fibroblast cells cultured on 2D with samples 3.5Ge-3.5Mu, 7Ge-7Mu, and 7Ge, 7 Mu hydrogels. (B) Quantification of percentage cell viability of fibroblasts for all 3.5Ge-3.5Mu, 7Ge-7Mu, and 7Ge, 7 Mu hydrogel samples. (C) Metabolic activity was assessed for 3.5Ge-3.5Mu, 7Ge-7Mu, and 7Ge, 7 Mu hydrogels. Student t-tests were performed for significant tests to compare each hydrogel composition between day 1 and day 3 (* *p* ≤ 0.05, *** *p* ≤ 0.001).

#### Liver cell-laden GelMA-MuMA hydrogel

GelMA-MuMA hybrid hydrogel was used to evaluate the potential of mucin based composite hydrogels in tissue engineering applications. Two different compositions of GelMA-MuMA hydrogels and GelMA alone were used to create cell-laden liver cancer tissue constructs using HepG2 cells. Cell encapsulation using all of these Muma hydrid gels show high viability for cells, thus confirming the high potential of GelMA-MuMA for microscale tissue engineering applications (**Fig. 7A**). The viability of the encapsulated cells was assessed on day 1 and day 3 (**Fig. 7B**). Cell proliferation at day 1 in all three hydrogels showed normalized fluorescence intensities of 2.1, 3.2, and 2.4 for 3.5Ge-3.5Mu, 7Ge alone, and 7Ge-7Mu, respectively. After three days of culture, a significant increase in cell proliferation was observed for 3.5Ge-3.5Mu and 7Ge alone, with 1.8- and 2-fold changes compared to day 1. In contrast, no significant change was observed for the 7Ge-7Mu hydrogel network after day 3, which confirms low cell proliferation in the GelMA-MuMA network with a total of 14 % solid content (p > 0.05). Cell viability has previously been shown to decrease with a higher percentage of solids (15%) in other hydrogel systems or GelMA alone (Nichol, Koshy et al. 2010). The primary reason behind the low viability of cells in a higher concentration of solids is higher stress due to stiffness arising from the network with smaller pore size, limiting the nutrient supply at the core, especially when cells are cultured for a longer time. Though both 3.5Ge-3.5Mu and 7Ge alone contained the same solid content, the hybrid hydrogels showed the lowest G1 value, the highest equilibrium swelling ratio, and a larger pore area (**Fig. 5A**). These properties of the hydrogel facilitate cell growth inside the 3D meshwork of the low-stiffness hydrogel. In addition, hybridization improved the cellular bioadhesion of MuMA based hydrogels (Sandberg 2009, Randall Raphael R. Janairo 2013).

**Fig 7.**
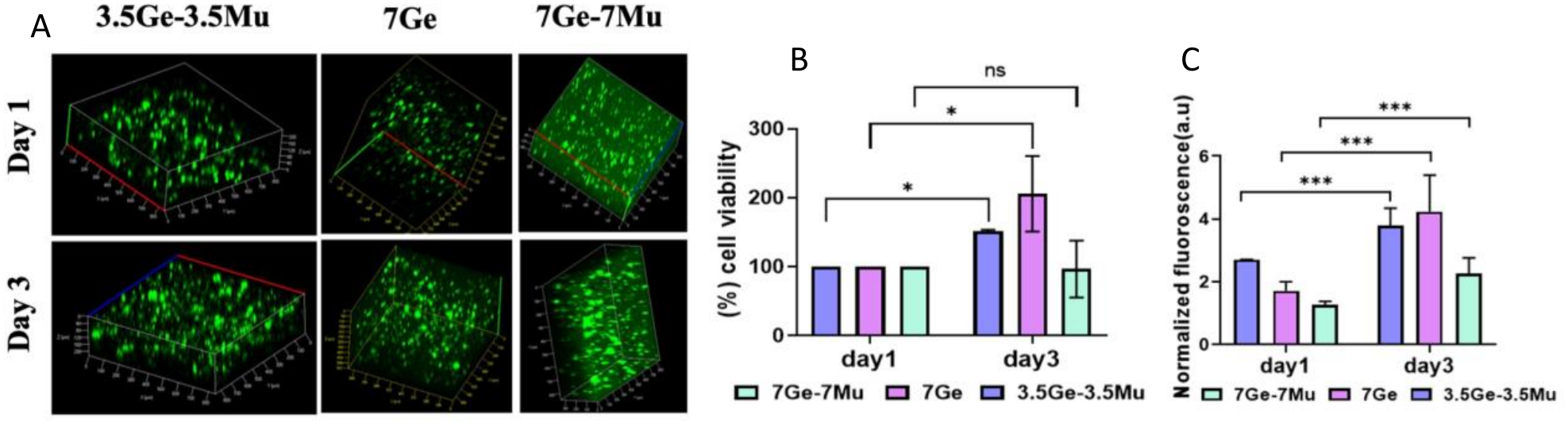
The calculated hydrogels percent (using equation 2) hemolysis compared to the positive control. **Lower panel:** Biocompatibility assay: (A) HepG2 cells encapsulated in 3.5Ge-3.5Mu, 7Ge-7Mu, and 7Ge hydrogels. Live cells were stained with calcein (green). (B) Cells in 3.5Ge-3.5Mu and 7Ge proliferated and showed excellent viability. (C) Metabolic activity was assessed for 3.5Ge-3.5Mu, 7Ge-7Mu, and 7Ge hydrogels. Student T-tests were performed for the significant tests to compare each hydrogel composition between day 1 and day 3 (* *p* ≤ 0.05, *** *p* ≤ 0.001).

It is worth mentioning that introducing GelMA facilitates cell encapsulation as GelMA contains an arginine-glycine-aspartic acid (RGD) peptide sequence, which promotes cell adhesion, proliferation, or differentiation. Moreover, the heavy side chains of gelatin possibly interact with MuMa to create interpenetrating networks, as evident from the thermal analysis data. Thus, the RGD peptide helps the cell spread in the hybrid hydrogel. On the other hand, MuMA helps achieve a low-stiffness hydrogel, which was challenging to obtain with either of the hydrogels alone below 5% (w/v). Cell metabolic activity was performed for cells encapsulated in the GelMA and MuMA networks during a similar time interval (**Fig. 7C**). Data showed a significantly higher increase in metabolic activity of 7% (w/v) hydrogels compared to 14% (w/v) hydrogels from day 1 to day 3. Specifically, an increase in normalized metabolic activity from 2.7 to 3.7, 1.7 to 4.2, and 1.3 to 2.3 was observed for the hydrogels: 3.5Ge-3.5Mu, 7Ge alone, and 7Ge-7Mu, respectively. These metabolic activity data were well-correlated with the cell proliferation results for all hydrogel compositions.

## CONCLUSIONS

In this study, MuMA was synthesized by methacrylation of mucin and subsequently crosslinked under UV irradiation to form hydrogels. MuMA hydrogels were prepared at different polymer concentrations, either alone or in combination with additives, and crosslinked for either 1 or 5 min. For hydrogels crosslinked for 5 min, physicochemical characterization revealed a clear relationship between MuMA concentration and hydrogel properties, whereas this correlation was less pronounced following 1 min of UV exposure.

In addition to polymer concentration and crosslinking time, hydrogel properties were further tuned through the incorporation of GelMA or glycerin. Hybrid GelMA-MuMA hydrogels were successfully formed using 3.5% (w/v) of each polymer, whereas MuMA alone failed to form stable hydrogels at concentrations below 5% (w/v). The incorporation of GelMA significantly altered the viscoelastic behavior of the hydrogels, resulting in a lower phase shift (Gʺ/Gʹ) compared with MuMA-only hydrogels, likely due to the formation of an interpenetrating polymer network. The hybrid hydrogels also exhibited larger pore sizes, slower degradation rates, and tunable mechanical properties that could not be achieved using either polymer alone, making them suitable for three-dimensional cell culture applications.

Glycerin acted as a plasticizer, increasing swelling while reducing the brittleness of MuMA hydrogel films and improving their flexibility. Thermal analysis confirmed interactions between MuMA and the incorporated additives, providing insight into the mechanisms underlying the observed changes in material properties. Furthermore, cells encapsulated within low-stiffness hydrogels exhibited enhanced proliferation and metabolic activity. The MuMA-based hydrogels demonstrated excellent biocompatibility and no detectable cytotoxicity, highlighting their potential for tissue engineering, disease modeling, cell encapsulation, and controlled drug delivery applications.

## STATISTICAL ANALYSIS

The data are presented as the means with standard error. One way ANOVA, or unpaired Student’s t-test, was used to compare differences as appropriate in the GraphPad Prism 5 software. Statistically significant differences were identified using a p-value lower than 0.05 as the cutoff value. Denotations for significance levels are defined in each figure *, p ≤ 0.05; **, p ≤ 0.01; ***, p < 0.001; ns, p > 0.05.

## SUPPORTING INFORMATION

The supplementary file is included.

## AUTHOR CONTRIBUTIONS

The manuscript is written by E.K, A.S.S and K.M and revised by E.K, A.S; C.U, R.H, M.J, S.K, N.F, F.Z, E.A, U.M, R.V, B.M, S.M and KM performed experiments. E.K, A.S.S, R.H, S.K; M.Z, B.M; S.M, R.B; and K.M interpreted the data. All authors have given approval to the final version of the manuscript.

## ACKNOWLEDGEMENT

This work was supported by Fulbright award and by the Deanship of Scientific Research (grants 2022-2021/198 and 2022-2021/199), University of Jordan-Amman-Jordan (Enam Khalil), by Terasaki Institute for Biomedical Innovation (TIBI). Authors would like to thank TIBI funding support and the National Institutes of Health (R01AR074234, R01GM126571, and U01CA214411). Authors would like to thank Prof. Ali Khademhosseini and Dr. Vadim Jucaud for reading and providing valuable feedback on the manuscript and Dr. Hossein Montazerian for his support in the rheology experiments.

## Data availability

(data transparency)

## Code availability

(software application or custom code)

## Authors’ contributions

(see Authorship principles section)

## ABBREVIATIONS

PEGTA: 4-arm poly(-ethylene glycol)-tetra-acrylate
RGD: arginine-glycine-aspartic acid
BSM: bovine submaxillary mucin
DiDW: deionized distilled water
DSC: Differential scanning calorimetry
DMSO: Dimethyl sulfoxide
DMEM: Dulbecco’s Modified Eagle Medium
ECM: extracellular matrix
FBS: Fetal Bovine Serum
GelMA: gelatin methacryloyl
Gly: glycerin
LVER: linear viscoelastic range
LAP: lithium phenyl-2,4,6-trimethylbenzoylphosphinate
MA: methacrylic anhydride
MuMA: mucin methacryloyl
NMR: Nuclear magnetic resonance
PBS: phosphate buffer saline
Tg: polymer glass transition temperature
PGM: porcine gastric mucin
RT: room temperature
SEM: scanning electron microscope
TGA: Thermal gravimetric analysis
VM: Vancomycin

