## Supplementary material for "Mucin methacryloyl (MuMA) based hydrogels: tunable features for tissue engineering and drug release control": NA

**Supplementary Information**

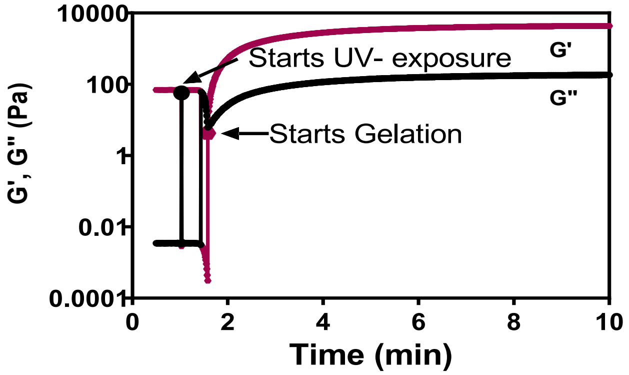


**Supplementary Figure 1:** Gelation kinetics of 10Mu hydrogel in PBS measured at 25 ˚C at 1 Hz frequency and 10% shear strain.

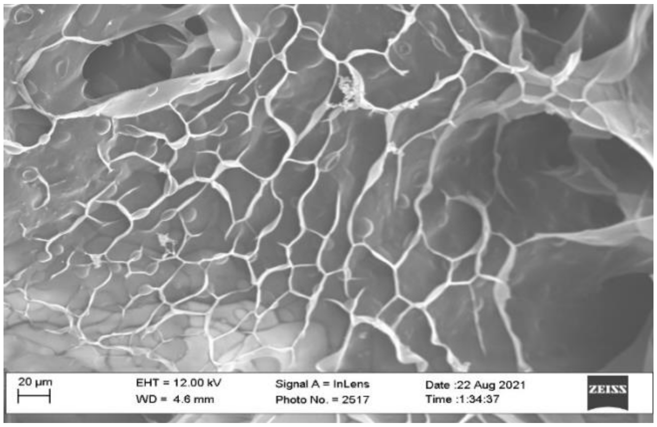


**Supplementary Figure 2**: SEM image of 3.5Ge-3.5Mu. scale bar equals 20 µm.

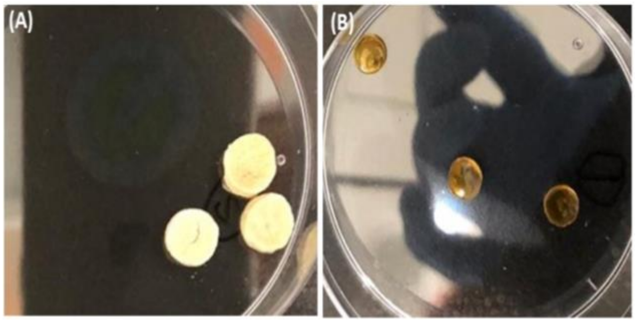


**Supplementary Figure 3. A**: 20Mu (freeze-dried) and **B:** 20Mu-20Gly (freeze-dried).

Hydrogel discs in both figures were prepared using 8 mm mold.
